# Elimination of Foxo triplets following glucocorticoid incubation dampened metabolic stress and enhanced anti-viral response targets based on transcriptomic analysis

**DOI:** 10.1101/2022.11.17.517006

**Authors:** Prasanth Puthanveetil, Daniel Rafferty, Boyi Gan, Charlotte A. Bolch

## Abstract

Glucocorticoids, both endogenous and exogenous, are known anti-inflammatory agents with associated adverse effects when administered chronically or in excess. Foxo family members, specifically Foxo1, Foxo3 and Foxo4 have demonstrated to be major regulators of metabolic complications following glucocorticoid action. We employed mouse embryonic fibroblasts as model system and using tamoxifen induced Cre+ recombinase activation, we achieved deletion of the Foxo triplets- Foxo1, Foxo3, and Foxo4. Following incubation of dexamethasone in both wild type and Foxo triplet deleted group, RNA sequencing was performed to reveal the transcriptome. Interestingly, the transcripts involved in initiating metabolic stress like PDK4 and 11 Beta HSD1 were downregulated, whereas transcripts involved in anti-viral response were upregulated following Foxo triplet deletion. The anti-inflammatory property of glucocorticoids was kept intact based on transcripts evaluated including GILZ and COX2. In summary, our study reveals a unique phenomenon where we will be able to manipulate glucocorticoid action by deleting Foxo triplets eliminating the metabolic stress components but enhancing anti-viral response and maintaining their clinical use as an anti-inflammatory agent. This work will lay foundation for a paradigm shift in clinical use of glucocorticoids according to therapeutic advantage with lesser adverse effects

## Introduction

Glucocorticoids (GC) as endogenous hormones are required not only during developmental stage but also for adult health and homeostasis [1,2]. In parity, exogenous glucocorticoids also have their own relevance in treating chronic inflammatory diseases like rheumatoid arthritis and airway diseases like asthma [3–7]. Problems arise when these endogenous secretions or exogenous supply become excess and unregulated leading to Cushing’s syndrome or Cushing’s like pathological states[7–15]. The notable features include weight gain accompanied by metabolic dysregulation and muscle atrophy and apparent metabolic disturbances accompanied by insulin resistance, hyperglycemia and hyperlipidemia[7–15]. Adrenalectomy has been shown to be an effective procedure in reducing the detrimental effects of glucocorticoid excess, but clinically it has various harmful aftermaths[16–18]. Previous findings from the applicant and other’s labs have demonstrated Foxo1 as a major regulator of glucocorticoid action[19,20].

Foxo1, is a transcription factor that has demonstrated transcriptional capabilities by binding to its own responsive elements[20–24]. At the same time in glucocorticoid mediated transcription function, Foxo1 has confirmed its role as a major transcription co-factor or co-regulator in enhancing PDK4 gene expression in cardiac and other cell types [20–24]. An enhanced PDK4 expression could lead to a reduction in glucose oxidation due to PDK4 mediated phosphorylation and inhibition of pyruvate dehydrogenase complex[20–24]. This stoppage of glucose metabolism could disturb the cellular metabolic homeostasis. On the other hand, FoxO1 has been shown to upregulate 11 β HSD1 in cell model of murine fibroblasts[25]. This role of FoxO1 was further validated with the help of a synthetic inhibitor of FoxO1, AS1842856 and insulin (a known natural cellular FoxO1 inhibitor), both of these agents were able to suppress the 11 β HSD1 gene expression[25]. With these observations, it suggests that FoxO family members not only regulate metabolic roles of GC action but also is involved in the generation of ligand via 11 β HSD1 for the glucocorticoid receptor.

Foxo1’s mandatory requirement as a key component in the glucocorticoid transcriptional machinery speaks of its eminence in glucocorticoid action. Along with FoxO1, other closely related Foxo family members; Foxo3 and Foxo4 have demonstrated to play this co-regulatory or co-transcriptional role in glucocorticoid mediated gene regulation[20,26–28]. Reports from Sprague-Dawley rat model of drug induced Cushing’s syndrome, where Cushing’s syndrome was induced by ACTH infusion using implanted osmotic pumps, demonstrated MuRF-1, and atrogin-1 gene increase in skeletal muscles[29]. This event was mediated by Foxo3a transcription factor. ACTH induced glucocorticoid action required the co-operation of Foxo3a transcription factor. Foxo3a also was demonstrated to be upregulated and a direct target of glucocorticoid receptor[29]. Interestingly in muscle cells of human (KM155C25 myotubes) and mouse (C2C12 myotube) origin and mouse muscle tissues, synthetic glucocorticoid, dexamethasone has been demonstrated to upregulate the Foxo triplets - Foxo1, Foxo3, and Foxo4 along with enhancing atrophy related genes; MuRF1 and MAFbx [30]. Will one Foxo family member compensate for the other or do they have specific co-regulatory role when it comes to targets regulated by glucocorticoids are not well established.

We have attempted to remove the triplet Foxo influence from GC mediated effect on transcription and for this we have made use of tamoxifen induced Cre+ recombinase technology to knock out the triplets-Foxo1/3/4. The derived Foxo1/3/4 L/L RosaCreERT2 mouse embryonic fibroblasts (MEFs) were either treated with vehicle (WT/controls) or tamoxifen to obtain the Foxo triple knock outs (TKO). Following Foxo triplet knock out, both WT/control and the TKO groups were subjected to dexamethasone exposure for short term. Our rationale was to obliterate the Foxo mediated effects under GC influence. Using this strategy, we were able to successfully disrupt the Foxo - GR association in GC excess mediated effects under glucocorticoid excess conditions. Our goal is to control or minimize the adverse metabolic consequences associated with glucocorticoid excess by deleting Foxo triplets out of the transcriptional machinery. By dissociating Foxo triplets from GC action we have mitigated metabolic stress but keeping the anti-inflammatory action of GCs intact. Our work will serve as stepping stone for researchers trying to curb the adverse effects due to endogenous and exogenous GC mediated effects either short term or chronically but maintaining the therapeutic effects of GCs intact.

## Materials and Methods

### Cell Culture Model

The cell culture model used for the study was Mouse Embryonic Fibroblasts (MEFs). MEFs were isolated from FoxO1/3/4, Rosa26-CreERT2 from E13.5 mouse embryos from a previously well established method[31–33]. These isolated cells were termed the group - WT cells. For deletion of Foxo triplets (FoxO1/3/4), we treat the WT cells in a DMEM with 4.5 g/l containing medium containing 10%FBS where 200 nM 4-hydroxy tamoxifen (4OHT) was incubated for at least 5 days along with undergoing passage. The required concentration of 4OHT was prepared by dissolving 4 OHT (sigma, Cat#: H7904,) by dissolving in appropriate volume of 100% EtOH. The 4 OHT mediated activation of Cre+ recombinase leads to Foxo triplet deletion. In the manuscript, this group is termed as Foxo triplet deleted group. Based on our prior and current experience[31,32], there is low expression of Cre+ as it is driven by an endogenous Rosa26 locus resulting in a decrease in knock out efficiency. We have chosen the dose and duration of treatment of 4 OHT based on keeping the balance between knock down efficiency and cell viability[31,32].

### Treatment groups

Before glucocorticoid treatment, high glucose (4.5 g/L) in 10 % FBS with antibiotics containing medium was substituted to a serum free low glucose (1 g/L glucose) containing medium with antibiotics. The goal is to avoid the high glucose and serum mediated cellular alteration in metabolism. WT and Foxo triplet KO MEFs were incubated with 100 nM dexamethasone for 4 hours in a 37 ^°^ C - CO_**2**_ containing incubator and then plated cells were removed and pure RNA extracted. This dose and time point was based on our previous work where dexamethasone demonstrated its gene transcriptional ability. Also based on our previous findings, early gene transcription function of glucocorticoids starts between 4-6 hours post ligand stimulation[20,34–36]. To identify direct gene targets and to avoid secondary transcriptional regulation early time point as 4 hours would be the most appropriate.

### Isolation of RNA

Following dexamethasone treatment of WT and Foxo triplet KO MEFs for 4 hours, RNA was isolated with PureLink^™^ RNA Mini Kit (Cat# 12183018A) using a previously well described protocol[37]. Once the pure RNA was extracted, we added RiboLock RNase Inhibitor (Cat # EO0381) at a concentration of 40 U/μL to prevent degradation of RNA. The RNA sample purity was determined using A260/280 and A260/230 values and samples with the desired purity was shipped for RNA sequencing (outsourcing at Omegabioservices, USA).

### RNA Library Preparation, Sequencing and bioinformatics based analysis

The RNA sequencing service was performed by Omegabioservices, GA, USA, an outsourcing service which provided the RNA library preparation, sequencing, and bioinformatics based analysis using a previously described and well established method[37].

### Statistical Analysis

Statistical Analysis was performed using Graph Pad Prism 9 software. The fold change values were calculated by Omegabioservices and expressed as log 2 fold change values of the obtained raw counts. The conversation to log 2 fold change was performed using DESeq2 method by our outsourcing vendor as described[37]. Comparisons of the mean values for the two groups were done using unpaired t-tests using Welch’s correction for unequal variances. The statistical significance was based on *p* values, *P<0.05* for all the evaluations. The comparison of the results expressed in raw values were represented as Mean ± SD. The log 2 fold change values were presented directly. The criteria for a significant change were based on both P-values of ***P<0.05*** and **log 2 fold** change value of ± 1.5 fold change.

## Results

### Knocking down the Foxo triplets in the presence of glucocorticoids reduce the expression of target genes regulating metabolic stress

To determine the role of Foxo triplets – Foxo1, 3 and 4 in glucocorticoid mediated transcription of known metabolic stress inducing gene targets-PDK4 and 11 beta HSD1, we deleted Foxo triplets in mouse embryonic fibroblasts (MEFs) using tamoxifen inducible Cre+ recombinase system using a well established method [31,32]. Following dexamethasone (100 nM) incubation for 4 hours, pure RNA samples were extracted and subjected to RNA sequencing. The known glucocorticoid receptor regulated targets were analyzed in WT and Cre+ induced MEFs following glucocorticoid stimulation. Evaluation of PDK4 and 11 beta HSD1 transcripts demonstrated a decrease of over 1.77 (>-1.77) and 2.02 (>-2.02) fold change as determined by a log 2-fold change values with a statistical significance of *P >0.05 (**Fig. 1 – left panel**)*. Along with these two targets we also analyzed some of the Foxo and glucocorticoid regulated common targets like PDK1, 2, 3, 11 beta HSD 2, Murf1 and Mafbx/Atrogin 1 (***Fig. 1 – right panel***). Even though the raw values for 11 beta HSD 2 following Foxo triplet deletion demonstrated statistical significance, the log 2 fold change value was only ***0.89-fold*** increase, and didn’t reach the minimum criterion of **+/-1.5 fold**. We did not observe any significant change in transcript levels for the other mentioned targets. These findings indicate that deletion of Foxo triplets during glucocorticoid stimulation decrease PDK4 and 11 beta HSD1 transcripts and could potentially attenuate the metabolic stress associated with PDK4 and 11 beta HSD1 (***Fig.1 – center panel***).

**Figure. 1.**
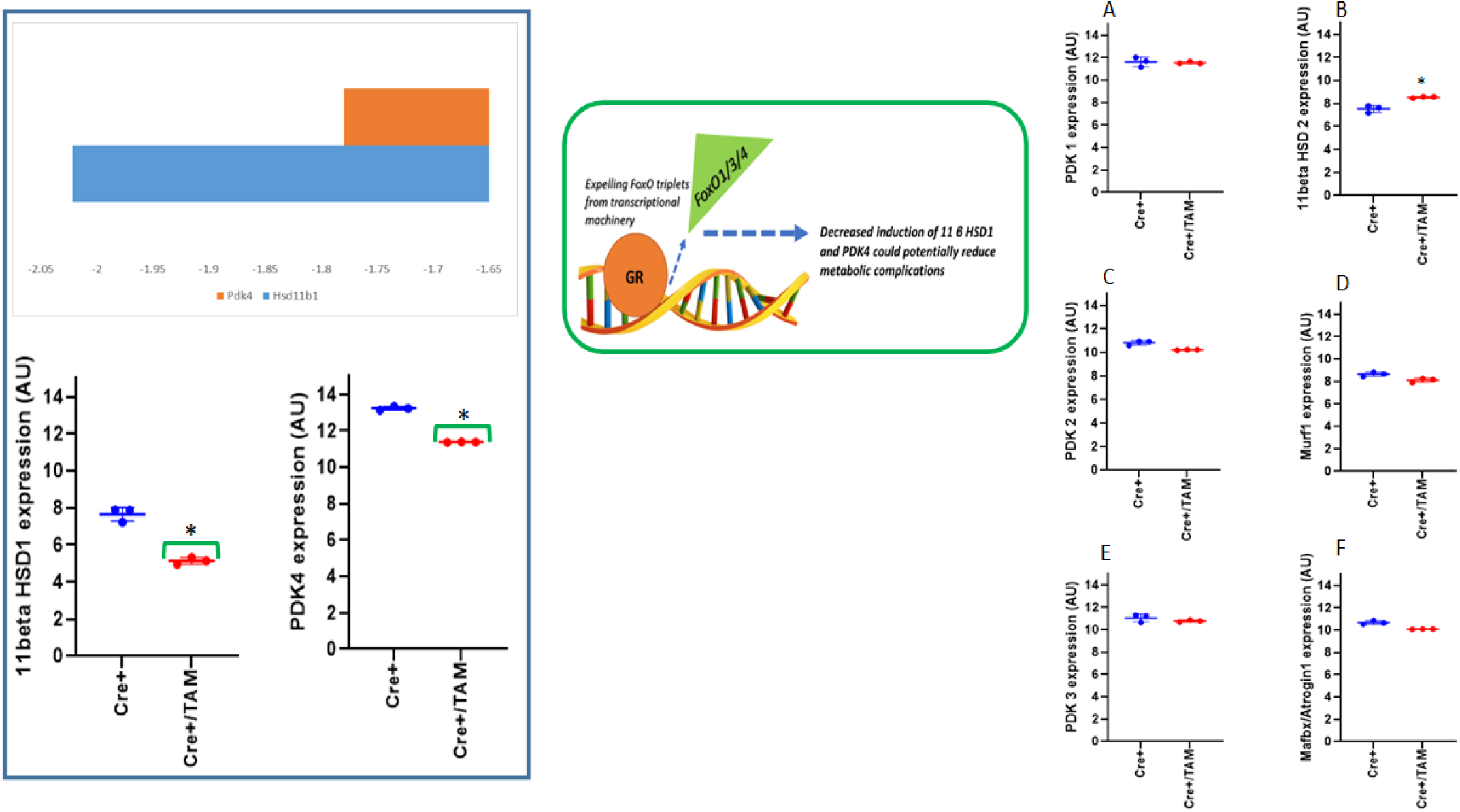
Regulation of transcripts involved in metabolic stress following Foxo triplet deletion in the presence of glucocorticoid incubation of MEFs. Dexamethasone incubation of WT and Triplet Foxo deleted MEFs for 4 hours demonstrated downregulation of transcripts for 11 beta HSD 1 and PDK4 transcripts (**left panel**). These metabolic stress inducing transcripts were downregulated > 1.5 fold (−1.5) as expressed by log 2 fold change values following Foxo triplet deletion in the presence of glucocorticoid treatment of MEFs (**center panel**). The other targets like PDK1 (**right panel-A**), PDK2 (**right panel-B**), PDK3 (**right panel-C**), 11 beta HSD 2 (**right panel-D**), Murf1 (**right panel-E**) and Mafbx/Atrogin 1 (**right panel-F**) did not the log 2 fold change values of ± 1.5 fold, even though the comparison of raw values for 11 beta HSD 2 demonstrated statistical significance. The statistics for determining significance was conducted using unpaired-t test followed by Welch’s correction in Graphpad Prism 9. The statistical significance, expressed as * ***P<0.05***.

### Deleting Foxo triplets following glucocorticoid stimulation does not interfere significantly with the anti-inflammatory response

GILZ has been demonstrated to be a direct target of glucocorticoid receptor which is involved in anti-inflammatory response[38]. We analyzed the regulation of GILZ and its known downstream targets[39,40]; PTGS2 (COX 2) and NOS 2 (iNOS) transcripts. Even though GILZ expression change demonstrated a significance of P<0.05, the log 2 fold change value observed was only <-0.774 (which was lower than our acceptance or set criteria of <-1.5 log 2 fold change value (***Fig.2A***). PTGS2 (COX 2) and NOS 2 (iNOS) transcripts did not change significantly (***Fig.2 B and C)***. Glucocorticoids are also known to interfere with NFκB transcription functiona[41] and down regulate inflammatory target genes like IL6[40], independent of GILZ. We evaluated the transcript levels of p65 subunit of NFκB (RelA - involved in active transcription) and also IL-6 (a major transcriptional target of NFκB). Interestingly neither Rel A (transcription component) nor its downstream inflammatory target, IL6 transcripts were affected by Foxo triplet down regulation (***Fig.2 D and E)***. The log 2 fold change values are stated in a chart and plotted using bar diagram (*(**Fig.2 – right lower panel***). Our data confirms that anti-inflammatory action of glucocorticoids are not regulated by Foxo triplets’ knock down (***Fig.2 - right upper panel***).

**Figure. 2.**
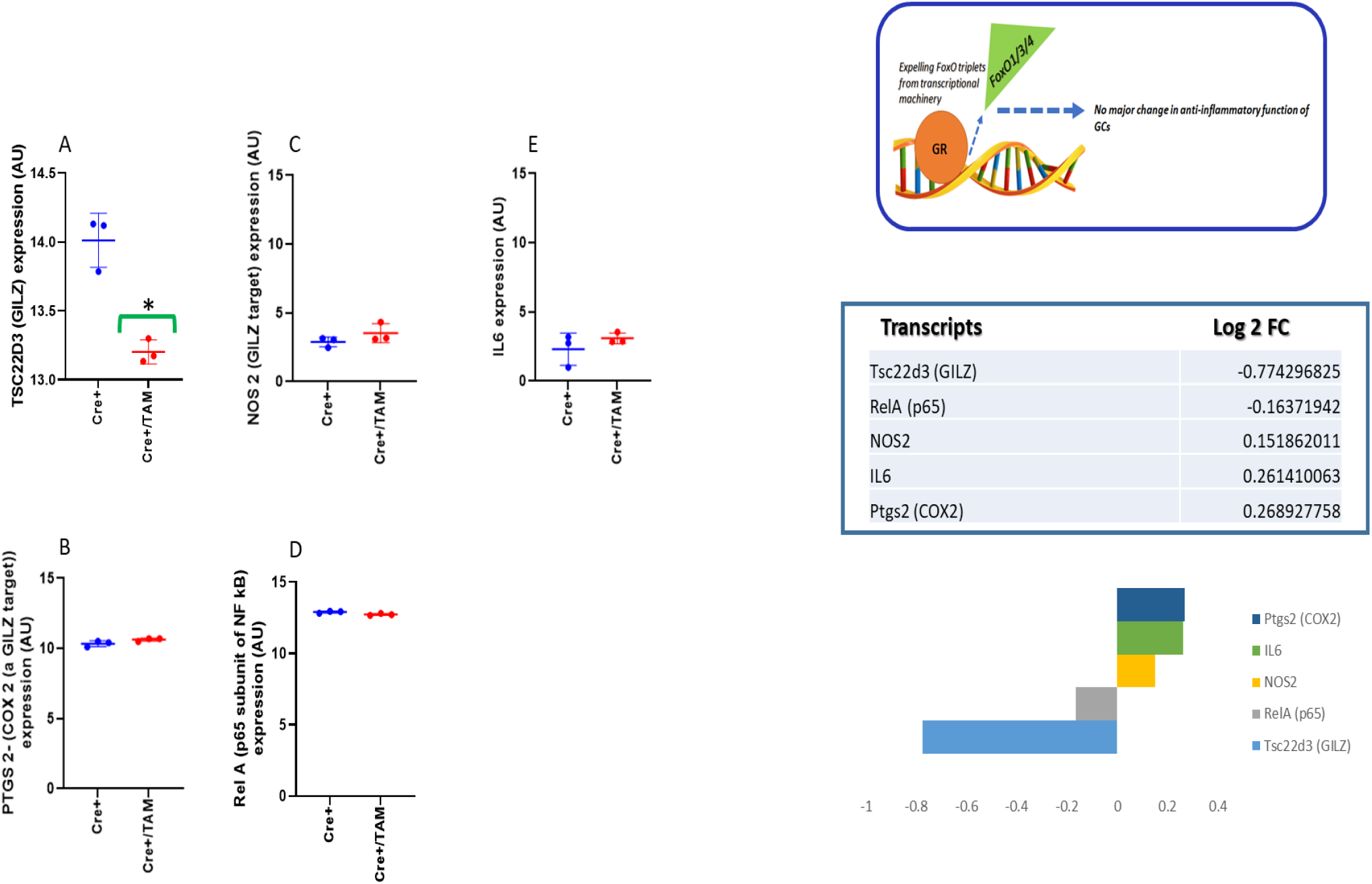
The impact of Foxo triplet deletion on anti-inflammatory targets of glucocorticoids. WT and Triplet Foxo deleted MEFs treated with dexamethasone demonstrated no major change in transcripts involved in regulating inflammation (**right-top panel**). Transcripts of Tsc22d3 (GILZ)-(**A)**, PTGS2 (COX2) (**B**), NOS 2 (**C**), Rel A (**D**) and IL 6 **(E)** did not demonstrate a log 2 fold change values of ± 1.5 fold, even though the Tsc22d3 (GILZ) raw values reached statistical significance. The log 2 fold change values for the transcripts which are glucocorticoid targets involved in inflammatory pathways are described in the chart and bar diagram (**right - middle and lower panels**). The raw values were compared using unpaired t-test with Welch’s correction and significance of ***P-value <0.05*** (*).

### Bioplanet based analysis revealed interferon, TGF beta and Oncostatin M signaling as the top regulated networks following Foxo triplets’ downregulation

Following RNA sequencing, Bioplanet based pathway analysis demonstrated that interferon signaling, TGF beta signaling and Oncostatin M signaling as the major gene networks regulated by Foxo triplet deletion in the presence of glucocorticoid treatment **(*Fig 3)***. Only these networks demonstrated statistical significance of *P<0.05 **(Fig 3 - insert).***

**Figure. 3.**
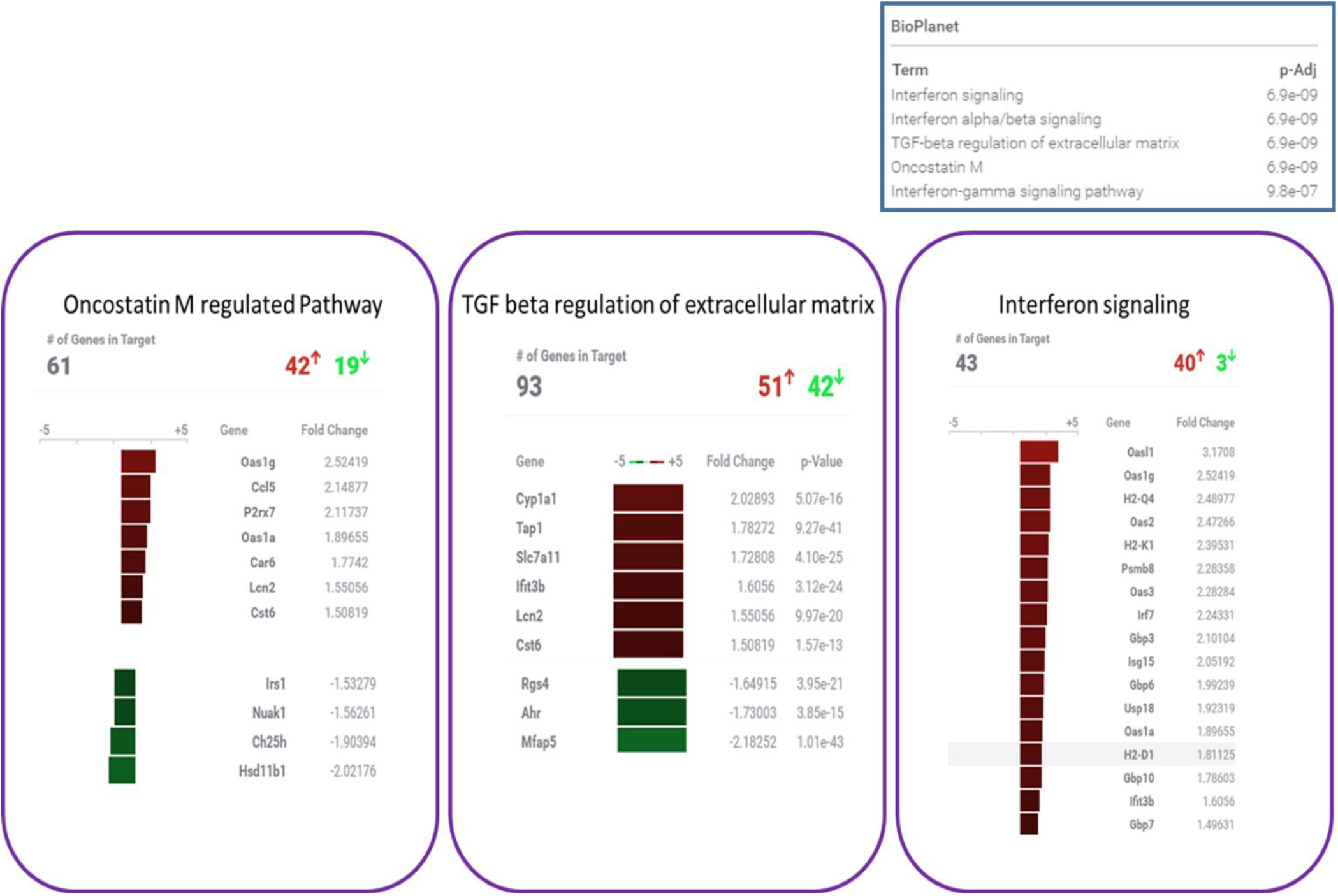
The pathways regulated by Foxo triplet deletion following glucocorticoid stimulation based on Bioplanet analysis. The differentially regulated transcripts following transcriptomic analysis were analyzed using Bioplanet based pathway analysis. Interferon signaling, TGF beta signaling, and Oncostatin M were the major networks regulated. Also, these were the only networks that demonstrated significance as per the adjusted P-values (**insert**).

### The top upregulated genes following Foxo triplets’ deletion exhibits involvement in anti-viral response

Following Foxo triplet deletion, the top upregulated genes were the ones involved in anti-viral response. Based on the log 2 fold change with significant increase, we noted that 2’5’ oligoadenylate synthetase-like 1 (Oasl1) was the top upregulated transcript (Ranked # 1) with > 3.17 fold change increase (***Fig.4A)***. Based on the descending order (ranking from 2 to 10), the upregulated transcripts; were Rsad2 (>2.99 fold change, ***Fig.4B***), Ifi44 (>2.94 fold change, ***Fig.4C***), Oas1g (>2.54 fold change, ***Fig.4D),*** H2-Q4 (>2.48 fold change, ***Fig.4E***), Oas2 (>2.47 fold change, ***Fig.4F***), CxCl10 (>2.43 fold change, ***Fig.4G***), ApoL9b (>2.40 fold change, ***Fig.4D***), H2-K1 (>2.39 fold change, ***Fig.4D***), and Fa2h (>2.38 fold change, ***Fig.4D***). Our data indicates that following triplet Foxo deletion, there is an upregulation of anti-viral response genes following glucocorticoid treatment **(*Fig.4 - right lower panel)***.

**Figure. 4.**
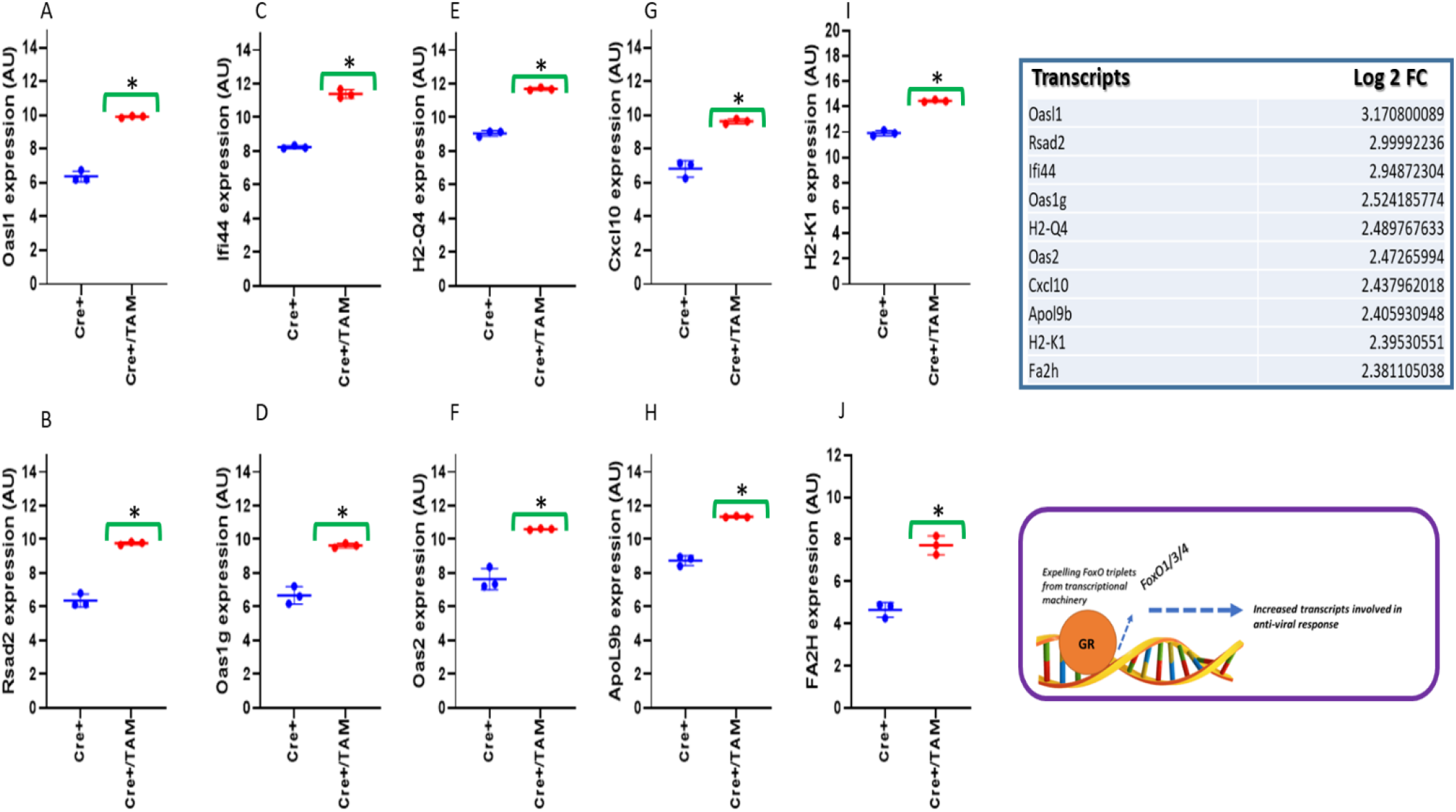
The top upregulated immune responsive transcripts following Foxo triplet deletion in presence of glucocorticoid treatment. Following Foxo triplet deletion, the transcripts that were mostly upregulated belonged to ones involved in immune responsiveness. The transcripts that were upregulated include (Oasl1) (***A)**,* Rsad2 (***B***), Ifi44 (***C***), Oas1g (***D),*** H2-Q4 **(*E***), Oas2 (***F**),* CxCl10 (***G***), ApoL9b (***D***), H2-K1 (***D***), and Fa2h (***D***). The log 2-fold changes values for the anti-viral transcripts are represented ***(right upper panel)*.** The mechanism depicting the effect of Foxo triplet deletion following glucocorticoid action with upregulation of gene involved in anti-viral response in represented (***right lower panel)***.

### Deletion of Foxo triplets does not influence glucocorticoid receptor expression

We were curious to know whether these alterations in target gene expressions we observe is due to alteration in glucocorticoid receptor expression. We analyzed the transcript levels of both glucocorticoid and mineralocorticoid receptors (***Fig.5A and B respectively***) and observed no significant change. This confirms that Foxo triplets’ deletion do not influence either glucocorticoid or mineralocorticoid receptor expression.

**Figure. 5.**
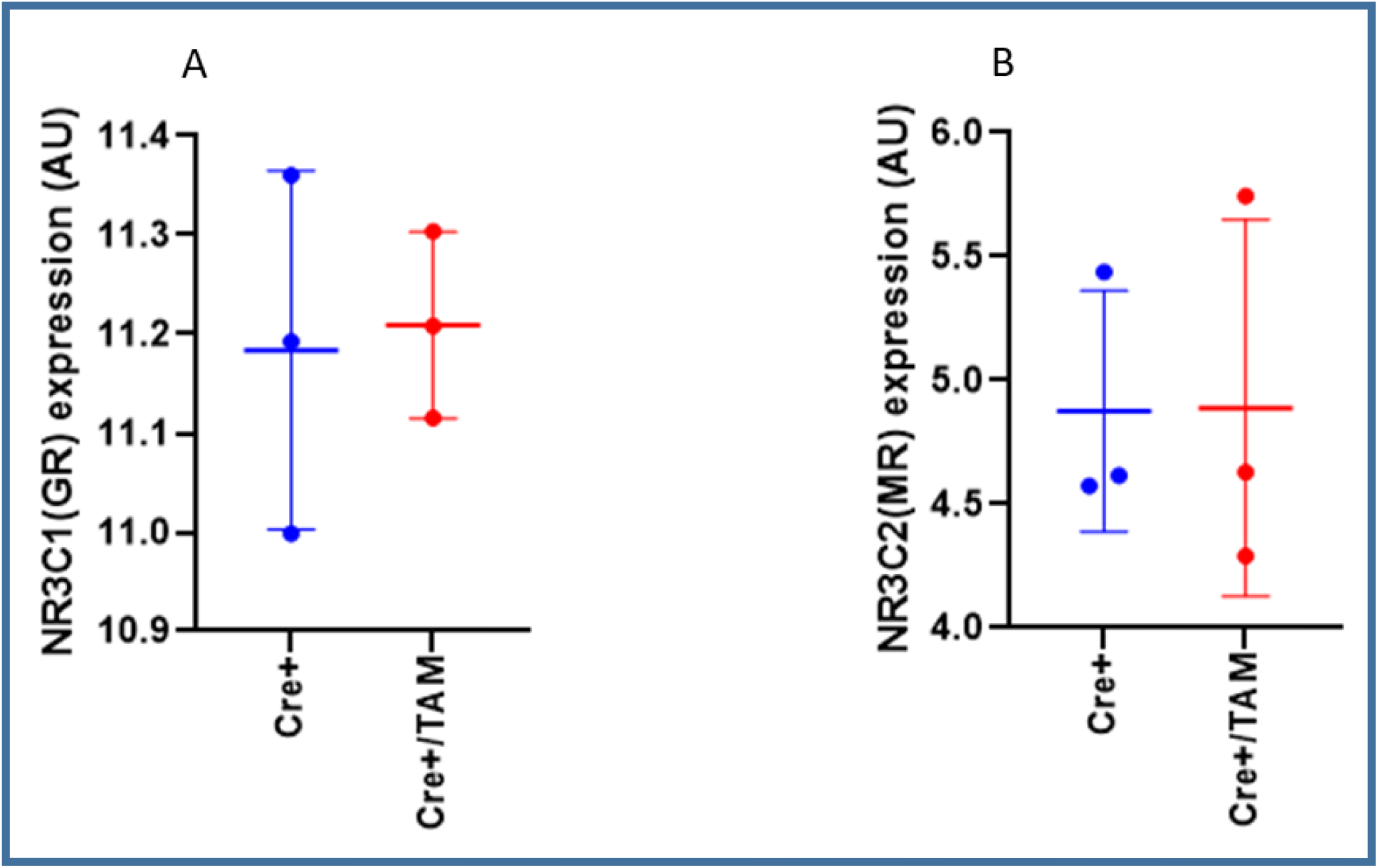
The effect of Foxo triplet depletion following glucocorticoid treatment on GR and MR transcripts: In the presence of glucocorticoid incubation, Foxo triplet deletion had no significant transcriptional effect on GR and MR genes (expressed in figures A and B) respectively.

## Discussion

Foxo family members, especially Foxo1, 3 and 4 have been reported to play an important role in this GC mediated metabolic alterations by acting as major nuclear transcriptional co-factor and target[20,26–29]. The targets regulated by Foxo under GC stimulated conditions like PDK4 and 11 beta HSD1 are also known regulators of metabolic stress[11,20]. When PDK4 brings about its action by inhibiting pyruvate dehydroganse complex activity and downregulating glucose oxidation, 11 beta HSD1 activity leads to enhanced bioconversion of inactive ligand to active endogenous ligand[11,20], which could cause undesired metabolic complications. Interestingly in our study following Foxo triplet knock down, in the presence of GC, the metabolic target genes, PDK4 and 11 beta HSD1 were downregulated by 1.77 and 2.02 as expressed by log 2-fold change values respectively. Downregulating 11 beta HSD1 could help in minimizing endogenous excess ligand generation and could prevent systemic metabolic complications due to cortisol excess as seen during Cushing’s syndrome and Cushing’s like conditions. Some of the adverse effects associated with excess cortisol have been linked to its nonspecific activation of mineralocorticoid receptor leading to hypertension, vasoconstriction, edema and fibrosis. By suppressing 11 beta HSD1 expression we would also benefit from excess MR activation. With PDK4 downregulation, the check on glucose oxidation could be removed and will be able to process glucose as substrate effectively to ATP. Inadvertently, this could lead to minimal lactate accumulation in metabolically active tissue, enhance glucose utilization and reduce the dependency on fatty acids, amino acids or even ketone bodies as substrate, maintaining the cellular metabolic homeostasis. The other isoforms of PDK including PDK1, PDK2, and PDK3 did not change significantly. This indicates that among the PDKs, PDK4 is the one regulated by Foxo’s in GC mediated transcription. The other isoform of 11 beta HSD1 enzyme is 11 beta HSD 2, involved in conversion of active form of GR ligand back to its inactive form, cortisone[42,43]. Unlike 11 beta HSD1 it’s counterpart 11 beta HSD2 did not change under Foxo triplet deletion. Role of Foxo family members especially, Foxo1, 3 and 4 have been linked to muscular atrophy by induction of Murf1 and Mafbx/Atrogin 1 under GC stimulation[26,29]. Following Foxo triplet downregulation, we did not observe any significant change in atrophy genes; Murf1 and Mafbx/Atrogin 1, which indicates that GCs don’t require Foxo family presence in regulating atrophy.

Along with regulating metabolism, GCs are also well-known as anti-inflammatory agents, whether they are endogenous or exogenous[28,39]. GCs bring about this function mostly through GILZ signaling nexus[28,39] or by suppressing NFκB mediated effect[41]. GCs by upregulating GILZ through a GR dependent mechanism suppress COX2 and iNOS induction, two major downstream targets of GILZ[28,39]. COX2 is the rate limiting enzyme for the induction of inflammatory eicosanoids including prostaglandins and leukotrienes[39]. iNOS is involved in generating peroxynitrite free radicals and creating an imbalance in cellular redox balance[39,40]. Interfering with NFκB mediated transcription, GCs are known to downregulate systemic inflammatory mediators like IL6[41]. In our study we analyzed whether Foxo triplets have any substantial contribution in the anti-inflammatory property of GC. We observed that even though a significant difference the log 2 fold change values only reached a decrease in 0.774 fold and did not fall under the inclusion criteria of ± 1.5 log 2 fold change. This minor change in GILZ was well translated into its downstream targets’ expression; both PTGS2 (COX 2) and NOS 2 (iNOS) transcripts did not demonstrate a change in log 2 fold change values and also statistical significance. The subunit of NFκB involved in transcriptional regulation; p65 (RelA) and its transcriptional target; IL-6 also did not exhibit any significant change. These observations have led us to derive the conclusion that by deleting Foxo triplets in the presence of GC treatment, the anti-inflammatory effects of GC are unaffected, and the targets are independent of Foxo triplet regulation.

Following RNA sequencing, BioPlanet platform based pathway analysis revealed that interferon signaling, TGF β signaling and Oncostatin M signaling are the major networks that were regulated when Foxo triplets were deleted following GC administration. Interferon signaling was ranked top in the list. Even though only 43 transcripts were regulated in this pathway, there were 16 transcripts which were significantly altered with ± 1.5 log 2-fold change. Interestingly all 16 transcripts were upregulated. Oncostatin M signaling had 61 transcripts altered but only 11 transcripts fulfilled the criteria of ± 1.5 log 2 fold change with statistical significance of P<0.05. Among the 11 transcripts changed significantly, 7 transcripts were upregulated and 4 downregulated. In TGF β signaling, even though 93 transcripts were altered, only 9 transcripts were changed significantly. Among those, 6 transcripts were upregulated and 3 were downregulated. Based on the number of significantly altered transcripts which are ± 1.5 log 2-fold change, interferon signaling has mounted up to the top ranking. Selection of top 10 regulated genes based on log 2 fold change values 9 out of 10 genes fell under the category of anti-viral and immune response genes. The top ranked in the list was Oasl1 which is encoded by the Oasl gene enhanced by > 3.17-fold change as expressed using log 2 fold change values. Oasl is known to demonstrate anti-viral response by preventing viral genome replication acting either through an IL-27 dependent or independent mechanism. This was followed by transcripts; Rsad2, Ifi44, Oas1g, H2-Q4, Oas2, CxCl10, ApoL9b, H2-K1, and Fa2h in that order. Unlike the top 9 genes, involved in anti-viral and immune response, Fa2h gene encodes for Fatty acid-2-hydroxylase protein. Fa2h is involved in the synthesis of 2-hydroxysphingolipids, the pre-cursor for the synthesis of spingolipids and glycospingolipids involved mostly in myelination or myelin sheath formation[44,45]. Overall analysis of this top 10 transcripts have revealed the fact that knocking out Foxo triples following GC administration could result in the enhancement of anti-viral and auto-immune response and prevention of neurodegeneration by its impact on myelination. Finally, we also evaluated whether knocking out Foxo triplets has got any role in regulating glucocorticoid receptor and its counterpart, mineralocorticoid receptor which endogenous GC and most synthetic GCs act as ligand. Interestingly we did not observe any change in GR and MR receptor expression levels. These findings reveal the fact that the alteration in transcriptome we observed is not due to alteration of expression of either GR or MR but Foxo triplets.

There are no studies till date which have investigated the contribution of Foxo triplets in glucocorticoid action. Unlike many other published works, the focus of this study is not to just re-establish a phenotype using a specific protein or proteins knockdown or knockin but rather to reveal a new pharmacological property of GC in the absence of Foxo triplets. Our study is the first to reveal the pharmacological mechanism that knocking down Foxo triplets in the presence of GCs could enhance i) anti-viral response ii) down regulation of metabolic stress genes and iii) maintaining the anti-inflammatory property of GCs (***Fig.6 – Summary***). Interestingly, this study reveals that by knocking down Foxo triplets in the presence of GCs we will be able to curtail the ligand generating enzyme 11 beta HSD1. This will be helping in curbing GC excess induced systemic complications as we see in Cushing’s syndrome and Cushing’s like metabolic complications. This could limit the positive feedback loop of excess GC induced metabolic complications. A unique property of GCs we observed during this study is that, by knocking down Foxo triplets we still maintain the anti-inflammatory property of GCs enhancing the anti-viral response through interferon pathway. For clinical practice, this work will provide insight into manipulating

**Figure. 6.**
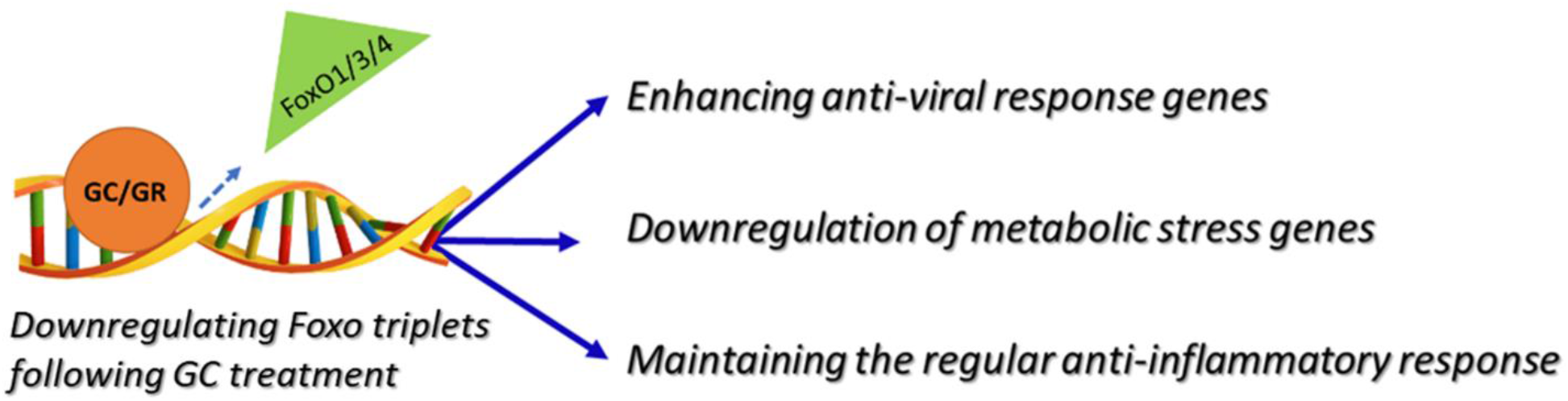
Summary diagram demonstrating the outcome of Foxo triplet downregulation following GC administration. Foxo triplet downregulation in the presence of glucocorticoid treatment short term leads to leads to transcripts involved in enhancing anti-viral response, downregulating metabolic stress but maintaining the inflammatory property.

GC actions, enhancing its beneficial effects just by regulating its co-factor/s involved in transcription, the Foxo triplets.

## Author contributions

PP conceived hypothesis, conducted experiments and collected data, wrote and edited the manuscript and was primarily involved in plotting the figures and making diagrams. All the authors have read and approved the manuscript. RFconducted experiments, and collected data. BG edited the manuscript and developed the cell line and contributed the cell line. CB edited the manuscript.

## Acknowledgements

I would like thank Dr. Mike Fay (Dean-CGS), and Dr. Philip Kopf (Chair-Department of Pharmacology), Dr. Walt Prozialeck, Dr. Joshua Edwards and Mr. Pete Lamar for sharing space and facilities and for their continuous support. We thank MWU core facility outsourcing funds and Office of Research Services for supporting this work.

## References

1. Gans, I.M.; Coffman, J.A. Glucocorticoid-Mediated Developmental Programming of Vertebrate Stress Responsivity. Front Physiol 2021, 12, 812195, doi:10.3389/fphys.2021.812195.

2. Quatrini, L.; Ugolini, S. New insights into the cell-and tissue-specificity of glucocorticoid actions. Cell Mol Immunol 2021, 18, 269–278, doi:10.1038/s41423-020-00526-2.

3. Mattishent, K.; Thavarajah, M.; Blanco, P.; Gilbert, D.; Wilson, A.M.; Loke, Y.K. Meta-review: adverse effects of inhaled corticosteroids relevant to older patients. Drugs 2014, 74, 539–547, doi:10.1007/s40265-014-0202-z.

4. Rossi, G.A.; Cerasoli, F.; Cazzola, M. Safety of inhaled corticosteroids: room for improvement. Pulm Pharmacol Ther 2007, 20, 23–35, doi:10.1016/j.pupt.2005.10.008.

5. Tilley, S.; Volmer, J.; Picher, M. Therapeutic applications. Subcell Biochem 2011, 55, 235–276, doi:10.1007/978-94-007-1217-1_9.

6. Walters, J.A.; Tan, D.J.; White, C.J.; Gibson, P.G.; Wood-Baker, R.; Walters, E.H. Systemic corticosteroids for acute exacerbations of chronic obstructive pulmonary disease. Cochrane Database Syst Rev 2014, CD001288, doi:10.1002/14651858.CD001288.pub4.

7. Weatherall, M.; Clay, J.; James, K.; Perrin, K.; Shirtcliffe, P.; Beasley, R. Dose-response relationship of inhaled corticosteroids and cataracts: a systematic review and meta-analysis. Respirology 2009, 14, 983–990, doi:10.1111/j.1440-1843.2009.01589.x.

8. Pivonello, R.; Faggiano, A.; Lombardi, G.; Colao, A. The metabolic syndrome and cardiovascular risk in Cushing’s syndrome. Endocrinol Metab Clin North Am 2005, 34, 327–339, viii, doi:10.1016/j.ecl.2005.01.010.

9. Fardet, L.; Feve, B. Systemic glucocorticoid therapy: a review of its metabolic and cardiovascular adverse events. Drugs 2014, 74, 1731–1745, doi:10.1007/s40265-014-0282-9.

10. Abraham, S.B.; Rubino, D.; Sinaii, N.; Ramsey, S.; Nieman, L.K. Cortisol, obesity, and the metabolic syndrome: a cross-sectional study of obese subjects and review of the literature. Obesity (Silver Spring) 2013, 21, E105–117, doi:10.1002/oby.20083.

11. Staab, C.A.; Maser, E. 11beta-Hydroxysteroid dehydrogenase type 1 is an important regulator at the interface of obesity and inflammation. J Steroid Biochem Mol Biol 2010, 119, 56–72, doi:10.1016/j.jsbmb.2009.12.013.

12. Paredes, S.; Ribeiro, L. Cortisol: the villain in metabolic syndrome? Rev Assoc Med Bras (1992) 2014, 60, 84–92, doi:10.1590/1806-9282.60.01.017.

13. Delaleu, J.; Destere, A.; Hachon, L.; Decleves, X.; Lloret-Linares, C. Glucocorticoids dosing in obese subjects: A systematic review. Therapie 2019, 74, 451–458, doi:10.1016/j.therap.2018.11.016.

14. Pivonello, R.; De Martino, M.C.; Iacuaniello, D.; Simeoli, C.; Muscogiuri, G.; Carlomagno, F.; De Leo, M.; Cozzolino, A.; Colao, A. Metabolic Alterations and Cardiovascular Outcomes of Cortisol Excess. Front Horm Res 2016, 46, 54–65, doi:10.1159/000443864.

15. Noppe, G.; van den Akker, E.L.; de Rijke, Y.B.; Koper, J.W.; Jaddoe, V.W.; van Rossum, E.F. Long-term glucocorticoid concentrations as a risk factor for childhood obesity and adverse body-fat distribution. Int J Obes (Lond) 2016, 40, 1503–1509, doi:10.1038/ijo.2016.113.

16. Liu, X.; Zhang, S.; Guo, Y.; Gang, X.; Wang, G. Treatment of Primary Pigmented Nodular Adrenocortical Disease. Horm Metab Res 2022, 54, 721–730, doi:10.1055/a-1948-6990.

17. Lase, I.; Gronberg, M.; Norlen, O.; Stalberg, P.; Welin, S.; Janson, E.T. Adrenalectomy in ectopic Cushing’s syndrome: A retrospective cohort study from a tertiary care centre. J Neuroendocrinol 2021, 33, e13030, doi:10.1111/jne.13030.

18. Kazi, S.D.; Andreone, M.A.; Radtke, L.E.; Strand, D.A.; Murphy, E.E. Cushing’s Syndrome in a Pregnant Patient Secondary to Adrenal Adenoma: Treatment With Unilateral Adrenalectomy. S D Med 2020, 73, 400–403.

19. Jiang, X.; Ma, H.; Li, C.; Cao, Y.; Wang, Y.; Zhang, Y.; Liu, Y. Effects of neonatal dexamethasone administration on cardiac recovery ability under ischemia-reperfusion in 24-wk-old rats. Pediatr Res 2016, 80, 128–135, doi:10.1038/pr.2016.54.

20. Puthanveetil, P.; Wang, Y.; Wang, F.; Kim, M.S.; Abrahani, A.; Rodrigues, B. The increase in cardiac pyruvate dehydrogenase kinase-4 after short-term dexamethasone is controlled by an Akt-p38-forkhead box other factor-1 signaling axis. Endocrinology 2010, 151, 2306–2318, doi:10.1210/en.2009-1072.

21. Kwon, H.S.; Huang, B.; Unterman, T.G.; Harris, R.A. Protein kinase B-alpha inhibits human pyruvate dehydrogenase kinase-4 gene induction by dexamethasone through inactivation of FOXO transcription factors. Diabetes 2004, 53, 899–910, doi:10.2337/diabetes.53.4.899.

22. Furuyama, T.; Kitayama, K.; Yamashita, H.; Mori, N. Forkhead transcription factor FOXO1 (FKHR)-dependent induction of PDK4 gene expression in skeletal muscle during energy deprivation. Biochem J 2003, 375, 365–371, doi:10.1042/BJ20030022.

23. Crossland, H.; Constantin-Teodosiu, D.; Greenhaff, P.L.; Gardiner, S.M. Low-dose dexamethasone prevents endotoxaemia-induced muscle protein loss and impairment of carbohydrate oxidation in rat skeletal muscle. J Physiol 2010, 588, 1333–1347, doi:10.1113/jphysiol.2009.183699.

24. Connaughton, S.; Chowdhury, F.; Attia, R.R.; Song, S.; Zhang, Y.; Elam, M.B.; Cook, G.A.; Park, E.A. Regulation of pyruvate dehydrogenase kinase isoform 4 (PDK4) gene expression by glucocorticoids and insulin. Mol Cell Endocrinol 2010, 315, 159–167, doi:10.1016/j.mce.2009.08.011.

25. Brazel, C.B.; Simon, J.C.; Tuckermann, J.P.; Saalbach, A. Inhibition of 11beta-HSD1 Expression by Insulin in Skin: Impact for Diabetic Wound Healing. J Clin Med 2020, 9, doi:10.3390/jcm9123878.

26. Senf, S.M.; Sandesara, P.B.; Reed, S.A.; Judge, A.R. p300 Acetyltransferase activity differentially regulates the localization and activity of the FOXO homologues in skeletal muscle. Am J Physiol Cell Physiol 2011, 300, C1490–1501, doi:10.1152/ajpcell.00255.2010.

27. Imae, M.; Fu, Z.; Yoshida, A.; Noguchi, T.; Kato, H. Nutritional and hormonal factors control the gene expression of FoxOs, the mammalian homologues of DAF-16. J Mol Endocrinol 2003, 30, 253–262, doi:10.1677/jme.0.0300253.

28. Latre de Late, P.; Pepin, A.; Assaf-Vandecasteele, H.; Espinasse, C.; Nicolas, V.; Asselin-Labat, M.L.; Bertoglio, J.; Pallardy, M.; Biola-Vidamment, A. Glucocorticoid-induced leucine zipper (GILZ) promotes the nuclear exclusion of FOXO3 in a Crm1-dependent manner. J Biol Chem 2010, 285, 5594–5605, doi:10.1074/jbc.M109.068346.

29. Kang, S.H.; Lee, H.A.; Kim, M.; Lee, E.; Sohn, U.D.; Kim, I. Forkhead box O3 plays a role in skeletal muscle atrophy through expression of E3 ubiquitin ligases MuRF-1 and atrogin-1 in Cushing’s syndrome. Am J Physiol Endocrinol Metab 2017, 312, E495–E507, doi:10.1152/ajpendo.00389.2016.

30. Cid-Diaz, T.; Santos-Zas, I.; Gonzalez-Sanchez, J.; Gurriaran-Rodriguez, U.; Mosteiro, C.S.; Casabiell, X.; Garcia-Caballero, T.; Mouly, V.; Pazos, Y.; Camina, J.P. Obestatin controls the ubiquitin-proteasome and autophagy-lysosome systems in glucocorticoid-induced muscle cell atrophy. J Cachexia Sarcopenia Muscle 2017, 8, 974–990, doi:10.1002/jcsm.12222.

31. Lin, A.; Yao, J.; Zhuang, L.; Wang, D.; Han, J.; Lam, E.W.; Network, T.R.; Gan, B. The FoxO-BNIP3 axis exerts a unique regulation of mTORC1 and cell survival under energy stress. Oncogene 2014, 33, 3183–3194, doi:10.1038/onc.2013.273.

32. Keniry, M.; Pires, M.M.; Mense, S.; Lefebvre, C.; Gan, B.; Justiano, K.; Lau, Y.K.; Hopkins, B.; Hodakoski, C.; Koujak, S.; et al. Survival factor NFIL3 restricts FOXO-induced gene expression in cancer. Genes Dev 2013, 27, 916–927, doi:10.1101/gad.214049.113.

33. Gan, B.; Lim, C.; Chu, G.; Hua, S.; Ding, Z.; Collins, M.; Hu, J.; Jiang, S.; Fletcher-Sananikone, E.; Zhuang, L.; et al. FoxOs enforce a progression checkpoint to constrain mTORC1-activated renal tumorigenesis. Cancer Cell 2010, 18, 472–484, doi:10.1016/j.ccr.2010.10.019.

34. Puthanveetil, P.; Wang, F.; Kewalramani, G.; Kim, M.S.; Hosseini-Beheshti, E.; Ng, N.; Lau, W.; Pulinilkunnil, T.; Allard, M.; Abrahani, A.; et al. Cardiac glycogen accumulation after dexamethasone is regulated by AMPK. Am J Physiol Heart Circ Physiol 2008, 295, H1753–1762, doi:10.1152/ajpheart.518.2008.

35. Kewalramani, G.; Puthanveetil, P.; Wang, F.; Kim, M.S.; Deppe, S.; Abrahani, A.; Luciani, D.S.; Johnson, J.D.; Rodrigues, B. AMP-activated protein kinase confers protection against TNF-{alpha}-induced cardiac cell death. Cardiovasc Res 2009, 84, 42–53, doi:10.1093/cvr/cvp166.

36. Kewalramani, G.; Puthanveetil, P.; Kim, M.S.; Wang, F.; Lee, V.; Hau, N.; Beheshti, E.; Ng, N.; Abrahani, A.; Rodrigues, B. Acute dexamethasone-induced increase in cardiac lipoprotein lipase requires activation of both Akt and stress kinases. Am J Physiol Endocrinol Metab 2008, 295, E137–147, doi:10.1152/ajpendo.00004.2008.

37. Puthanveetil, P.; Kong, X.; Brase, S.; Voros, G.; Peer, W.A. Transcriptome analysis of two structurally related flavonoids; Apigenin and Chrysin revealed hypocholesterolemic and ketogenic effects in mouse embryonic fibroblasts. Eur J Pharmacol 2021, 893, 173804, doi:10.1016/j.ejphar.2020.173804.

38. Wang, J.C.; Derynck, M.K.; Nonaka, D.F.; Khodabakhsh, D.B.; Haqq, C.; Yamamoto, K.R. Chromatin immunoprecipitation (ChIP) scanning identifies primary glucocorticoid receptor target genes. Proc Natl Acad Sci U S A 2004, 101, 15603–15608, doi:10.1073/pnas.0407008101.

39. Yang, N.; Zhang, W.; Shi, X.M. Glucocorticoid-induced leucine zipper (GILZ) mediates glucocorticoid action and inhibits inflammatory cytokine-induced COX-2 expression. J Cell Biochem 2008, 103, 1760–1771, doi:10.1002/jcb.21562.

40. Vago, J.P.; Galvao, I.; Negreiros-Lima, G.L.; Teixeira, L.C.R.; Lima, K.M.; Sugimoto, M.A.; Moreira, I.Z.; Jones, S.A.; Lang, T.; Riccardi, C.; et al. Glucocorticoid-induced leucine zipper modulates macrophage polarization and apoptotic cell clearance. Pharmacol Res 2020, 158, 104842, doi:10.1016/j.phrs.2020.104842.

41. Auphan, N.; DiDonato, J.A.; Rosette, C.; Helmberg, A.; Karin, M. Immunosuppression by glucocorticoids: inhibition of NF-kappa B activity through induction of I kappa B synthesis. Science 1995, 270, 286–290, doi:10.1126/science.270.5234.286.

42. Kucka, M.; Vagnerova, K.; Klusonova, P.; Miksik, I.; Pacha, J. Corticosterone metabolism in chicken tissues: evidence for tissue-specific distribution of steroid dehydrogenases. Gen Comp Endocrinol 2006, 147, 377–383, doi:10.1016/j.ygcen.2006.02.007.

43. Christy, C.; Hadoke, P.W.; Paterson, J.M.; Mullins, J.J.; Seckl, J.R.; Walker, B.R. 11beta-hydroxysteroid dehydrogenase type 2 in mouse aorta: localization and influence on response to glucocorticoids. Hypertension 2003, 42, 580–587, doi:10.1161/01.HYP.0000088855.06598.5B.

44. Maldonado, E.N.; Alderson, N.L.; Monje, P.V.; Wood, P.M.; Hama, H. FA2H is responsible for the formation of 2-hydroxy galactolipids in peripheral nervous system myelin. J Lipid Res 2008, 49, 153–161, doi:10.1194/jlr.M700400-JLR200.

45. Kawaguchi, M.; Sassa, T.; Kidokoro, H.; Nakata, T.; Kato, K.; Muramatsu, H.; Okuno, Y.; Yamamoto, H.; Kaname, T.; Kihara, A.; et al. Novel biallelic FA2H mutations in a Japanese boy with fatty acid hydroxylase-associated neurodegeneration. Brain Dev 2020, 42, 217–221, doi:10.1016/j.braindev.2019.11.006.

